# Stomatal density affects gas diffusion and CO_2_ assimilation dynamics in Arabidopsis under fluctuating light

**DOI:** 10.1101/2020.02.20.958603

**Authors:** Kazuma Sakoda, Wataru Yamori, Tomoo Shimada, Shigeo S. Sugano, Ikuko Hara-Nishimura, Yu Tanaka

**Author notes:** Corresponding Author: Kazuma Sakoda, Graduate School of Agriculture, Kyoto University, Kitashirakawa Oiwake-cho, Sakyo-ku, Kyoto 606-8502, Japan. **Abbreviations:** *A*, CO_2_ assimilation rate; *CAR*, cumulative CO_2_ assimilation rate; *CER*, cumulative transpiration rate; *C*_i_, intercellular CO_2_ concentration; *E*, transpiration rate; *gs*, stomatal conductance; *L*_g_, guard cell length; PPFD, photosynthetic photon flux density; *SD*, stomatal density; *WUE*, water use efficiency; *WUE*_i_, integrated water use efficiency.

## Abstract

Stomatal density (*SD*) is closely associated with photosynthetic and growth characteristics in plants. In the field, light intensity can fluctuate drastically throughout a day. The objective of the present study is to examine how the change in *SD* affects stomatal conductance (*gs*) and CO_2_ assimilation rate (*A*) dynamics, biomass production, and water use under fluctuating light. Here, we compared the photosynthetic and growth characteristics under constant and fluctuating light among four lines of *Arabidopsis thaliana* (L.): a wild-type (WT), a *STOMAGEN/EPFL9*-overexpressing line, *STOMAGEN/EPFL9*-silencing line, and an *EPIDERMAL PATTERNING FACTOR 1* knockout line (*epf1*). Lower *SD* resulted in faster response of *A* owing to the faster response of *gs* to fluctuating light and higher water use efficiency without decreasing *A*. Higher *SD* resulted in a faster response of *A* because of the higher initial *gs*. *epf1*, with a moderate increase in *SD*, showed the larger carbon gain, attributable to the high capacity and fast response of *A*, yielding higher biomass production than WT under fluctuating light. The present study suggests that higher *SD* can be beneficial to improve biomass production in the plant under field conditions.

## Introduction

Stomata, pores on the epidermis of plant leaves, function to maintain the balance between CO_2_ uptake for photosynthesis and water loss for transpiration (Mcadam and Brodribb, 2012). The regulation of stomatal development has an important role in plant adaption to long-term environmental changes, and stomatal differentiation and patterning are tightly regulated by various signals influenced by plant hormones and environmental stimuli (Qi and Torii, 2018). The peptide signals in a family of EPIDERMAL PATTERNING FACTOR (EPF) were identified to function in the stomatal development of Arabidopsis (*Arabidopsis thaliana* (L.) Heynh) (Hara et al., 2007). It is demonstrated that EPF1 and EPF2 combine with the receptor-like protein, TOO MANY MOUTHS (TMM) and ERECTA family leucine-rich repeat-receptor-like kinases and, consequently, restrain a specific process in stomatal development. Contrastingly, STOMAGEN/EPFL9 combines with TMM competitively to EPF1 and EPF2, and promote stomatal development (Lee et al., 2015; Sugano et al., 2010). Elucidation of the genetic factors involved in stomatal development has enabled us to modify the stomatal size, number, and patterning in plant leaves by transgenic approach.

Enhancing leaf photosynthesis has been attempted to drive further increase in biomass production in crop plants (von Caemmerer and Evans, 2010; Sakoda *et al*., 2016; Yamori *et al*., 2016b). Gas diffusional resistance from the atmosphere to the chloroplast is one of the limiting factors for leaf photosynthetic capacity (Farquhar and Sharkey, 1982). Furthermore, it has been highlighted that the conductance to gas diffusion via stomata (*gs*) can be a major determinant of CO_2_ assimilation rate (*A*) (Wong et al., 1979). The potential of *gs* is mainly determined by the size, depth, and opening of single stoma, and their density (Franks and Beerling, 2009). It has been controversial how the change in the stomatal density (*SD*), defined as the stomata number per unit leaf area, affects photosynthetic and growth characteristics in plants (Lawson and Blatt, 2014). Doheny-Adams *et al*., (2012) reported that lower *SD* yielded higher growth rate and biomass production in Arabidopsis under constant light owing to the favorable water condition and temperature for metabolism and low metabolic cost for stomatal development (Doheny-Adams et al., 2012). Contrastingly, lower *SD* resulted in the depression of *gs* and/or *A* in Arabidopsis and poplar plants (Bussis *et al*., 2006; Yoo *et al*., 2010; Wang *et al*., 2016). An *SDD1* knockout line of Arabidopsis with higher *SD* showed higher *gs* and *A* than a wild-type line, depending on light condition (Schlüter et al., 2003). Previously, we reported that higher *SD* by overexpressing *STOMAGEN/EPFL9* resulted in the enhancement of *gs* and *A* in Arabidopsis under constant and saturated light conditions (Tanaka et al., 2013). Therefore, *SD* manipulation could have the potential to enhance photosynthetic and growth characteristics in plants, even though that effect can depend on the species or environmental conditions.

In the field, light intensity can fluctuate at different scales, from a few seconds to minutes, over the course of a day owing to changes in the solar radiation, cloud cover, or self-shading in the plant canopy. The gradual increase in *A* can be shown after the transition from low to high light intensity, and this phenomenon is called “photosynthetic induction.” A simulation analysis demonstrated that the potential loss of the cumulative amount of CO_2_ assimilation caused by photosynthetic induction can reach at least 21% in wheat (*Triticum aestivum* L.) and soybean (*Glycine max* (L.) Merr.) (Tanaka et al., 2019; Taylor and Long, 2017). In rice (*Oryza sativa* L.) and soybean, there is genotypic variation in the speed of photosynthetic induction, which causes significant differences in the cumulative carbon gain under fluctuating light (Adachi et al., 2019; Soleh et al., 2017, 2016). Consequently, speeding up photosynthetic induction can yield more efficient carbon gain, which will open a new pathway to improve biomass production in plants under field conditions.

Photosynthetic induction is typically limited by three phases of the biochemical and diffusional processes: (1) activation of electron transport, (2) activation of the enzymes in the Calvin-Benson cycle, and (3) stomatal opening (Pearcy, 1990; Yamori, 2016a). Especially, the activation of Rubisco (5–10 min for full induction) and stomatal opening (20–30 min for full induction) constitute a major limitation to photosynthetic induction (Carmo-Silva and Salvucci, 2013; Yamori et al., 2012). The overexpression of *PATROL1*, controlling the translocation of a major H^+^-ATPase (AHA1) to the plasma membrane, resulted in faster *gs* response to fluctuating light in Arabidopsis without the change in *SD* (Hashimoto-Sugimoto et al., 2013). Arabidopsis knockout mutants of ABA transporter, which plays pivotal roles in stomatal closure, improved stomatal response to fluctuating light and photosynthesis (Shimadzu et al., 2019). Furthermore, the rapid movement of stomata is important for plants to achieve high water use efficiency (*WUE*) (Qu et al., 2016). Notably, the overexpression of *BLINK1*, light-gated K^+^ channel in guard cells, accelerates the stomatal opening and closing, which improved biomass production without additional water loss in Arabidopsis under the fluctuating light (Papanatsiou et al., 2019). These facts evidence that rapid stomatal movement can be beneficial for the effective carbon gain and water use under fluctuating light conditions. However, how *SD* changes affect *gs* and *A* dynamics, biomass production, and water use under these conditions has been understudied (Drake et al., 2013; Papanatsiou et al., 2016; Schuler et al., 2018).

The objective of this study was to examine how the change in *SD* affects the photosynthetic and growth characteristics in plants under fluctuating light conditions. Therefore, we investigated the dynamics of *g*s, *A*, and transpiration rate (*E*) by gas exchange measurements and biomass production under fluctuating light conditions in four Arabidopsis lines differing in *SD*. The results shine a light on the manipulation of *SD* to improve the biomass production and water use in plants under field.

## Results

### Stomatal density, size, and clustering

Compared with WT, ST-OX showed 268.1% higher *SD* (*p* < 0.05) while ST-RNAi showed 70.9% lower *SD* (*p* < 0.05) as reported in Sugano *et al*. (2010) and Tanaka *et al*. (2013) (Fig. 1A). *L*_g_ of ST-OX was 10.0% lower than that of WT (*p* < 0.01) (Fig. 1B). *SD* in *epf1* was 46.5% higher than that of WT (*p* < 0.1) and *L*_g_ of *epf1* was slightly higher than that of WT as reported in Dow. *et al*. (2014) (*p* < 0.1). Stomatal clustering was not observed in WT and ST-RNAi, while two to five stomata were clustered in ST-OX and *epf1* (Fig. 1C). The ratio of stomata in each clustering category was higher in ST-OX than that in *epf1*.

**Fig. 1.**
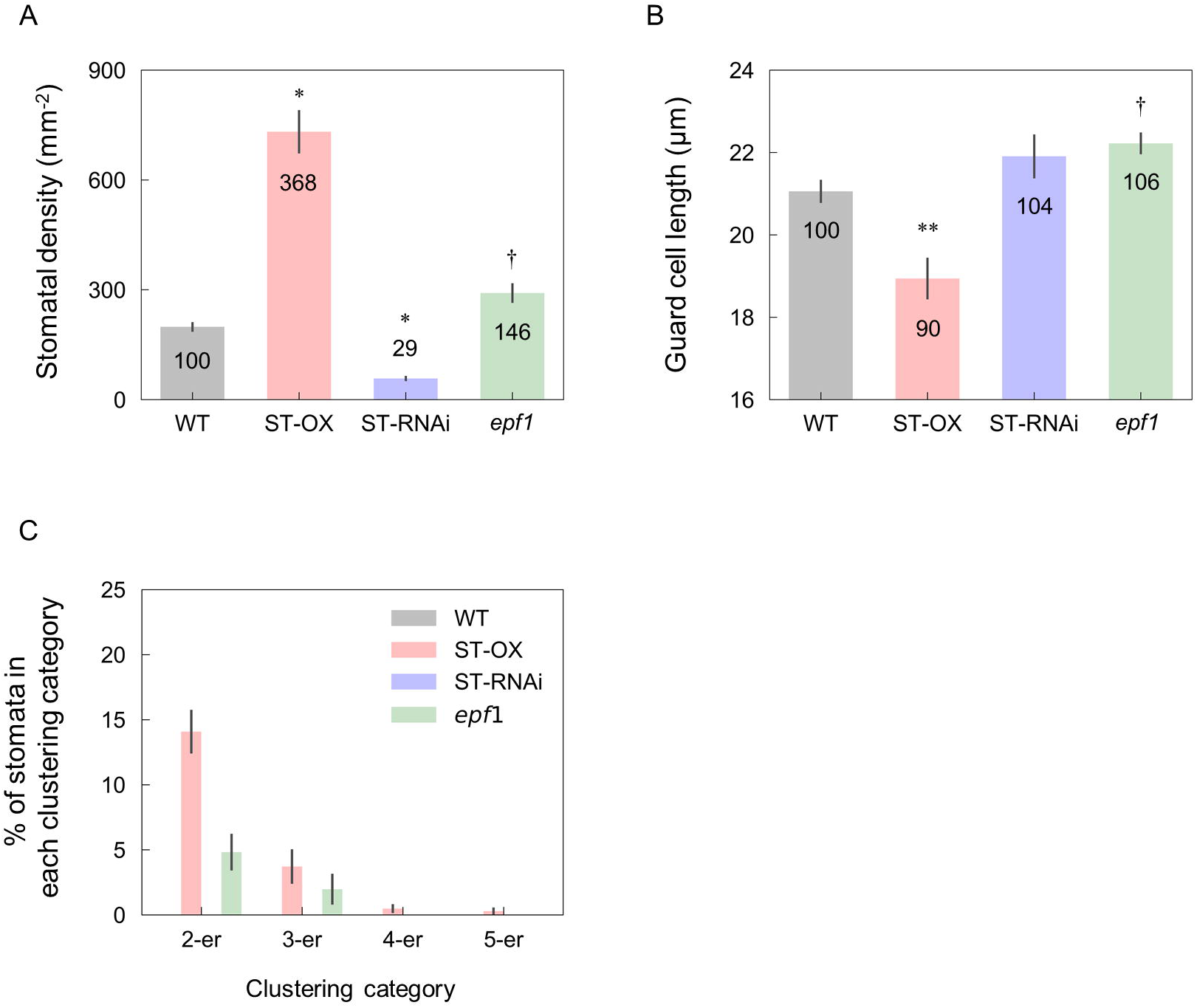
Stomatal density, size, and clustering. (A) The stomatal density (*SD*), (B) guard cell length (*L*_g_) and (C) the rate of stomata in 2–5 clustering category were evaluated on the fully expanded leaf in a wild-type line (WT), a *STOMAGEN*/*EPFL9* overexpressing (ST-OX) or amiRNA-mediated silencing line (ST-RNAi), and an *EPF1* knockout line (*epf1*) of *Arabidopsis thaliana*. The vertical bars indicate the standard error (n = 6). †, * and ** indicate the significant variation in each parameter between WT and each transgenic line at *p* < 0.1, 0.05, and 0.01, respectively, according to the Steel test in (A) or Dunnett’s test in (B) and (C). The values in each column represent the relative value of each line to WT.

### Photosynthetic characteristic under fluctuating light conditions

ST-OX and *epf1* maintained higher *gs*, *A*, and *E* than WT under fluctuating light condition (Fig. 2B, D, and E). ST-OX showed higher *C*_i_ and lower *WUE* than WT throughout the measurement period, while *epf1* only showed higher *C*_i_ and lower *WUE* than WT during the initial 50−60 min after the dark condition (Fig. 2C and F). ST-RNAi showed higher *gs*, *C*_i_, *A*, and *E* than WT in the initial induction phase and, subsequently, similar or lower values of these parameters than those of WT. *WUE* of ST-RNAi was constantly higher than that of WT except for the initial induction phase.

**Fig. 2.**
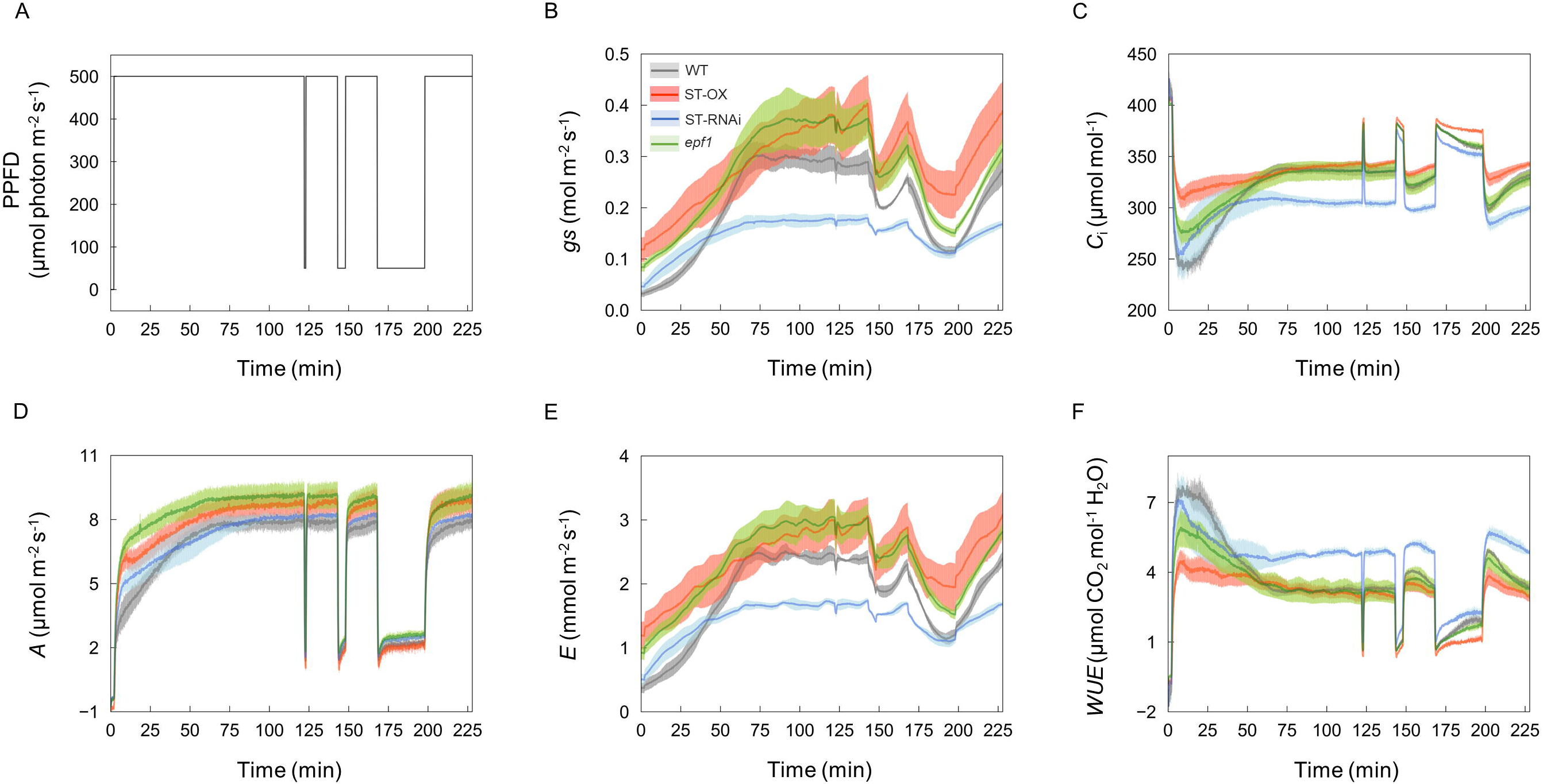
Photosynthetic characteristics under the non-steady state. (A) Light intensity set-up as for (B) stomatal conductance (*gs*), (C) intercellular CO_2_ concentration (*C*_i_), (D) CO_2_ assimilation rate (*A*), (E) transpiration rate (*E*), and (F) water use efficiency (*WUE*) measurements on the fully expanded leaf in the four lines of Arabidopsis. The gas exchange measurements were conducted at a CO_2_ concentration of 400 ppm and air temperature of 25°C. Vertical bars indicate the standard error (n = 3).

After the light intensity change from dark to high (500 μmol photon m^-2^ s^-1^), *gs* response was faster in ST-RNAi and slower in ST-OX than that in WT (Fig. 3A). In ST-OX, *gs* did not reach the steady state even after 120 min with illumination (Fig. 2B). *t*_dark_*gs*0.6_ in ST-RNAi was smaller (*p* < 0.05) and *t*_dark_*gs*0.9_ in ST-OX was larger than those in WT (*p* < 0.05) (Fig. 3B and C). *gs* response in *epf1* was similar to that in WT. The response of *A* in all transgenic lines was faster than that in WT after light intensity change from dark to high intensity (Fig. 3D). *t*_dark_*A*0.6_ in ST-OX (*p* < 0.05), ST-RNAi (*p* < 0.1), and *epf1* (*p* < 0.05) were shorter than that in WT (Fig. 3E and F). There was no significant variation in *t*_dark_*A*0.9_ between WT and each line, although *t*_dark_*A*0.9_ in *epf1* was 37.0% shorter than that of WT.

**Fig. 3.**
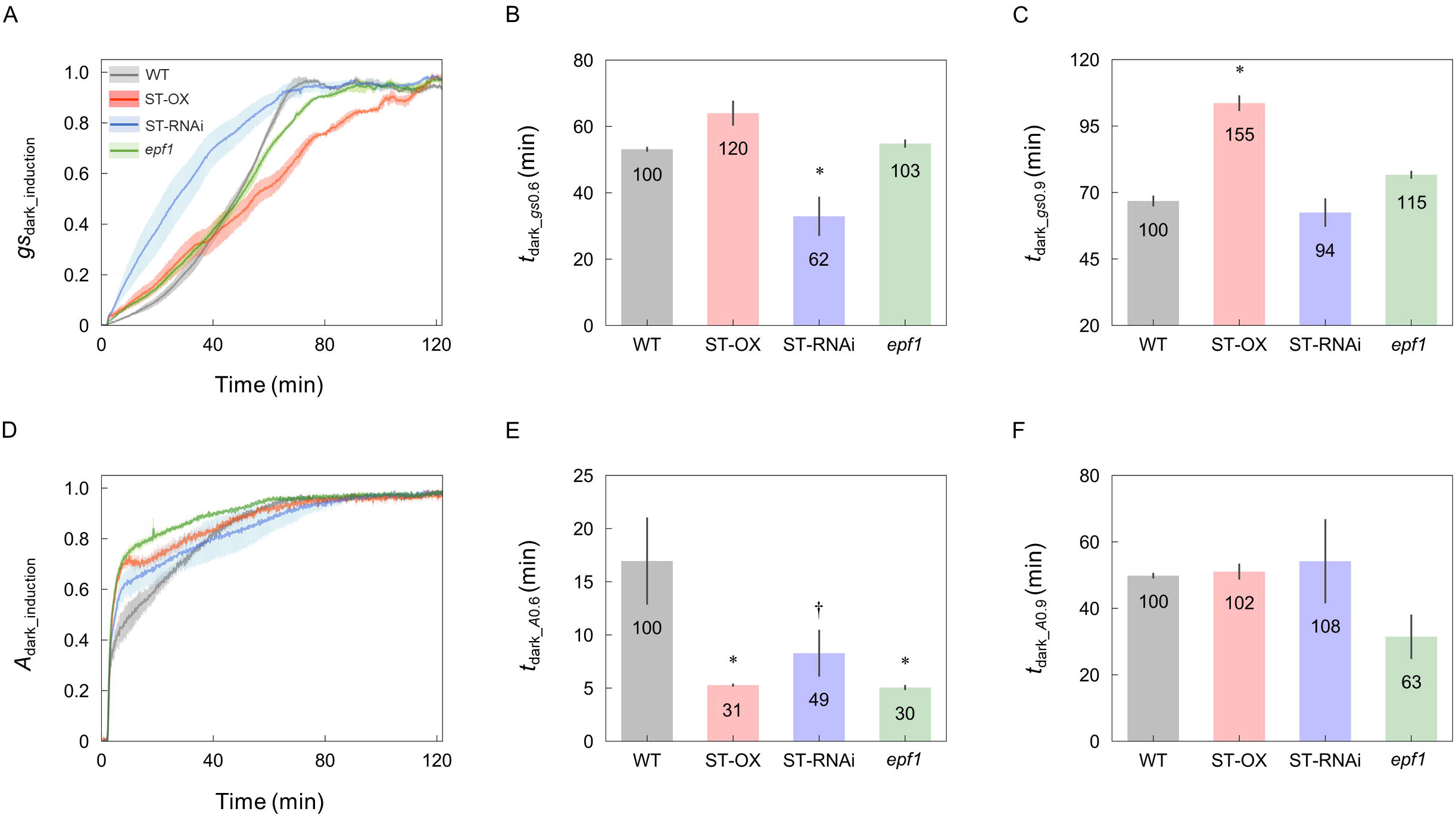
Time lapsed until reaching 0.6 and 0.9, out of a maximum value of 1, for stomatal conductance and CO_2_ assimilation rate increase after changing light intensity from dark to high. The induction state of (A) stomatal conductance (*gs*) and (D) CO_2_ assimilation rate (*A*) was evaluated in the four lines of Arabidopsis based on *gs*_dark_induction_ and *A*_dark_induction_ defined as eqn. 1 and 2, respectively, under a PPFD of 500 µmol photon m^-2^ s^-1^ for 120 min after the dark period. The time when (B, C) *gs*_dark_induction_ and (E, F) *A*_dark_induction_ reached 0.6 (*t*_dark_*gs*0.6,_ *t*_dark_*A*0.6_) and 0.9 (*t*_dark_*gs*0.9,_ *t*_dark_*A*0.9_), respectively, was compared between WT and each transgenic line. Vertical bars indicate the standard error (n = 3). † and * indicate significant differences in each parameter between WT and each transgenic line at *p* < 0.1 and 0.05, respectively, according to Dunnett’s test. The values in each column represent the relative value of each line to WT.

After light intensity changed from low (50 μmol photon m^-2^ s^-1^) to high (500 μmol photon m^-2^ s^-1^), a large variation in *gs* and *A* responses was not shown between WT and each transgenic line (Fig. 4A and C). There was no significant variation in *t*_low_*gs*0.6_ between WT and each transgenic line (Fig. 4B). Although *gs* response in ST-OX seemed faster than that in WT (Fig. 4A), *gs*_low*_*induction_ was overestimated in ST-OX because *gs* did not reach the steady state in 120 min with illumination after the dark period (Fig. 2B and 4A). *t*_low_*gs*0.9_ was not calculated since *gs* did not reach 0.9 to *gs*_max_ in any lines except for ST-OX. *t*_low_*A*0.6_ was significantly shorter in three transgenic lines than that in WT (*p* < 0.05), while the variation was quite small (Fig. 4D). *t*_low_*A*0.9_ in ST-OX was 52.8% shorter than that in WT (*p* < 0.1) (Fig. 4E).

**Fig. 4.**
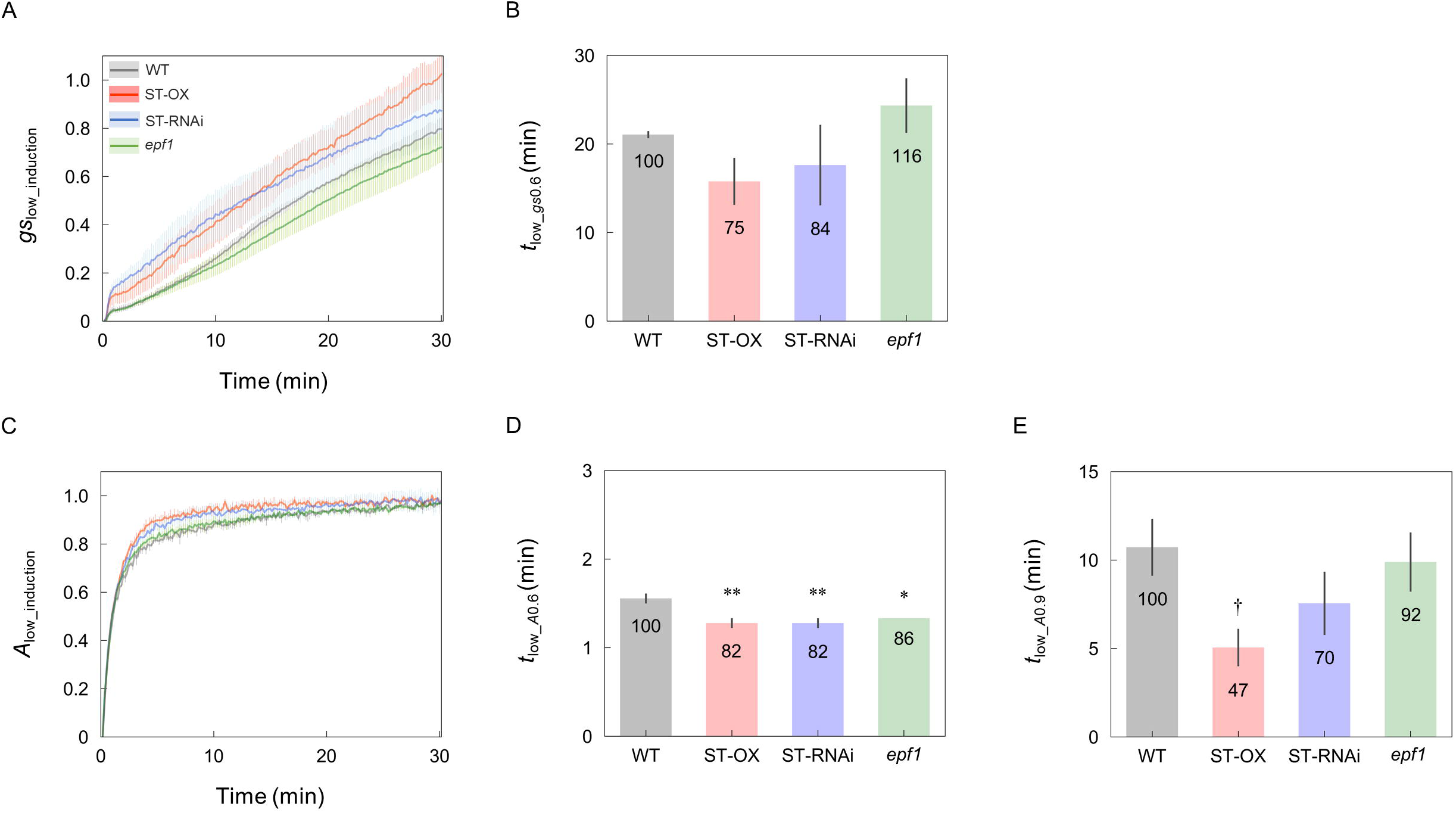
Time lapsed until reaching 0.6 and 0.9, out of a maximum value of 1, for stomatal conductance and CO_2_ assimilation rate increase after changing light intensity from low to high. The induction state of (A) stomatal conductance (*gs*) and (C) CO_2_ assimilation rate (*A*) was evaluated in the four lines of Arabidopsis based on *gs*_low_induction_ and *A*_low_induction_ defined as eqn. 3 and 4, respectively, under a PPFD of 500 µmol photon m^-2^ s^-1^ for 30 min after a PPFD of 50 µmol photon m^-2^ s^-1^ for 30 min. The time when (B) *gs*_low_induction_ and (D, E) *A*_low_induction_ reached 0.6 (*t*_low_*gs*0.6,_ *t*_low_*A*0.6_) and 0.9 (*t*_low_*A*0.9_) was compared between WT and each transgenic line. Vertical bars indicate the standard error (n = 3). †, *, and ** indicate significant differences in each parameter between WT and each transgenic line at *p* < 0.1, 0.05, and 0.01, respectively, according to Dunnett’s test. The values in each column represent the relative value of each line to WT.

In the steady state under dark conditions, *gs*_dark_ in ST-OX and *epf1* were 264.5% (*p* < 0.01) and 160.6% (*p* < 0.1) higher, respectively, than that in WT (Fig. 5A). *t*_dark_*A*0.6_ decreased with the increase in *gs*_dark_ when *gs*_dark_ < 0.055, and it was constantly independent of *gs*_dark_ for *gs*_dark_ > 0.055 (Fig. 5B). *t*_dark_*A*0.9_ showed no relation to *gs*_dark_ (Fig. 5C). *gs*_low_ in ST-OX was 95.6% higher than that in WT (*p* < 0.01) (Fig. 5D). *t*_low_*A*0.6_ and *t*_low_*A*0.9_ were independent of *gs*_low_ (Fig. 5E and F). Under the steady state at 500 μmol photon m^-2^ s^-1^, there was no significant difference in *gs*, *A*, and *E* between WT and ST-OX or *epf1*, although the transgenic lines showed values of these parameters higher than those in WT (Fig. S1). ST-RNA_i_ showed lower *C*_i_ (*p* < 0.05), higher *WUE* (*p* < 0.05), and similar *A* than WT. *gs* and *E* in ST-RNAi tended to be lower than those in WT although there was no significant variation.

**Fig. 5.**
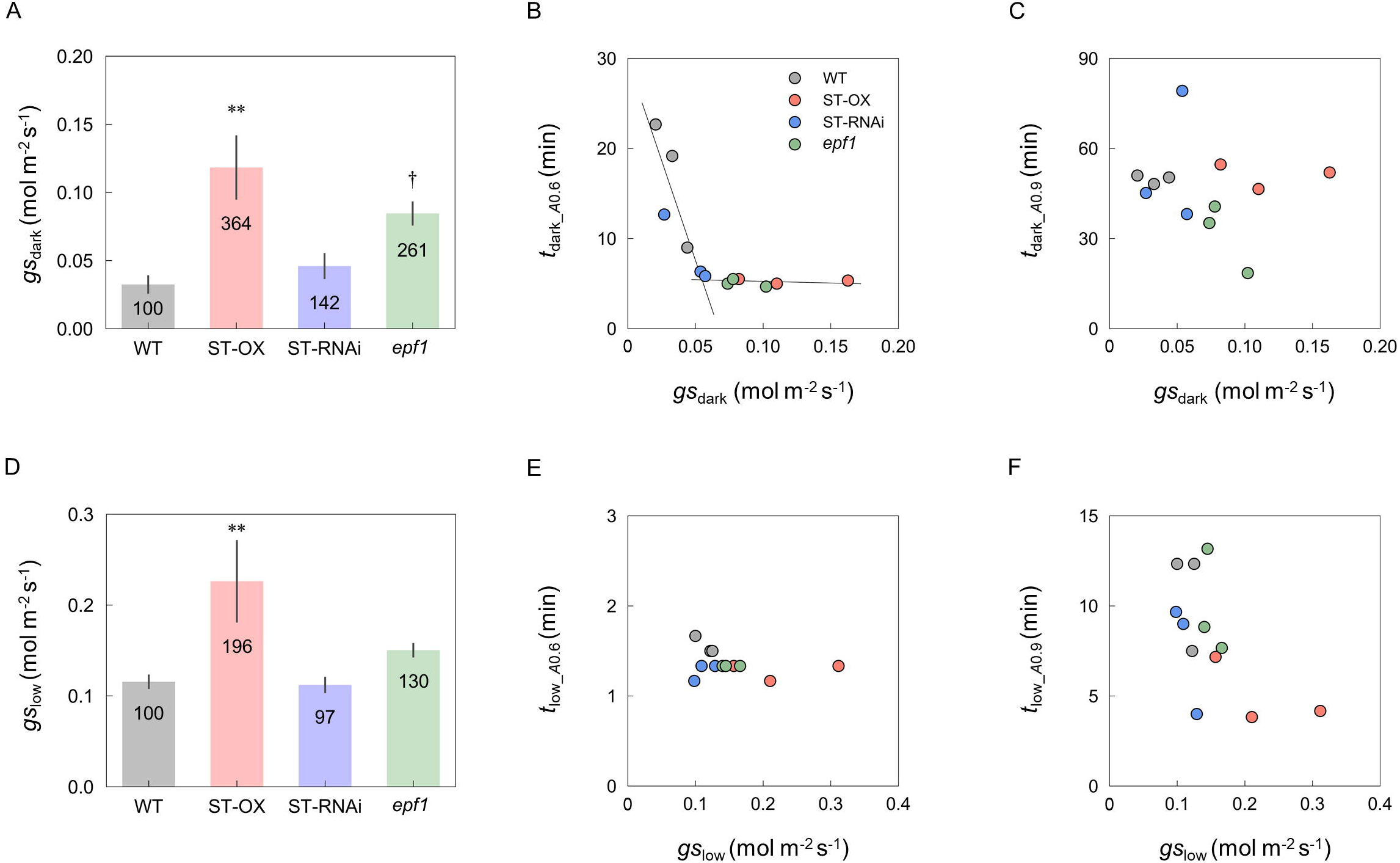
Relationship between the initial stomatal conductance and time lapsed until reaching 0.6 and 0.9, out of a maximum value of 1, for CO_2_ assimilation rate increase after changing light intensity. The steady-state value of stomatal conductance under (A) the dark condition (*gs*_dark_) and a PPFD of 50 µmol m^-2^ s^-1^ (*gs*_low_) were compared between WT and each transgenic line. The relationship was investigated between *gs*_dark_ or *gs*_low_ and the time to reach (B, E) 0.6 (*t*_dark_*A*0.6_, *t*_low_*A*0.6_) or (C, F) 0.9 (*t*_dark_*A*0.9_, *t*_low_*A*0.9_), out of a maximum value of 1, for CO_2_ assimilation rate increase after changing the light intensity from the dark or low to high. Vertical bars indicate the standard error (n = 3). †and ** indicate significant differences in each parameter between WT and each transgenic line at *p* < 0.1 and 0.01, respectively, according to Dunnett’s test. The values in each column represent the relative value of each line to WT.

*CAR* in *epf1* was 20.6% higher than that in WT (*p* < 0.1) (Fig. 6A). The loss rate in *epf1* was 33.4% lower than that in WT, although the variation was not significant (Fig. 6B). *CER* in ST-OX and *epf1* were 30.6% and 27.5% higher, respectively, than that of WT with no significant variation (Fig. 6C). ST-RNAi showed similar *CAR* to that of WT, while it showed 21.0% lower *CER*, which contributed to *WUE*_i_ being 32.1% higher than that of WT (*p* < 0.1) (Fig. 6D).

**Fig. 6.**
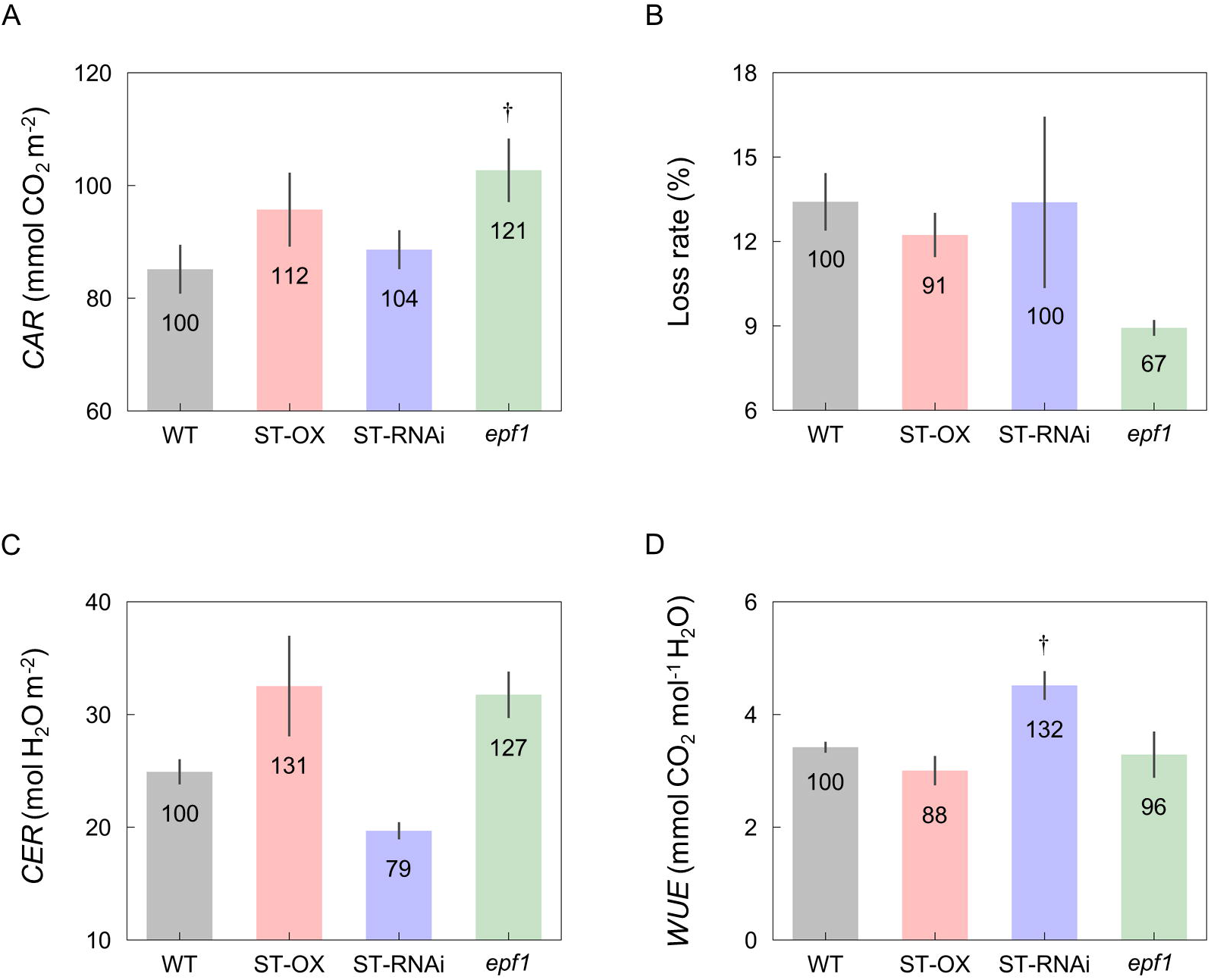
Cumulative carbon gain and water use under fluctuating light. (A) Cumulative rate of CO_2_ assimilation (*CAR*) and (B) transpiration (*CER*) was measured in the four lines of Arabidopsis under the fluctuating light described in Fig. 2A. (C) The loss rate of CO_2_ assimilation caused by the induction response was calculated based on eqn. 3. (D) Integrated water use efficiency (*WUE*_i_) was calculated as the ratio of *CAR* to *CER*. Vertical bars indicate the standard error (n = 3). † indicates significant differences in each trait between WT and each transgenic line at *p* < 0.1 according to Dunnett’s test. The values in each column represent the relative value of each transgenic line to WT.

### Biomass production under constant and fluctuating light conditions

Compared with WT, dry weight of the above ground biomass under constant light (*DW*_constant_) of *epf1* was similar, while that under fluctuating light (*DW*_fluctuating_) of *epf1* was 25.6% higher than that of WT (*p* < 0.05) (Fig. 7C and D). *DW*_constant_ and *DW*_fluctuating_ of ST-OX were slightly lower or higher than those of WT, respectively, with no significance variation. *DW*_constant_ and *DW*_fluctuating_ of ST-RNAi were 44.9% and 36.5% lower than those of WT, respectively (*p* < 0.01).

**Fig. 7.**
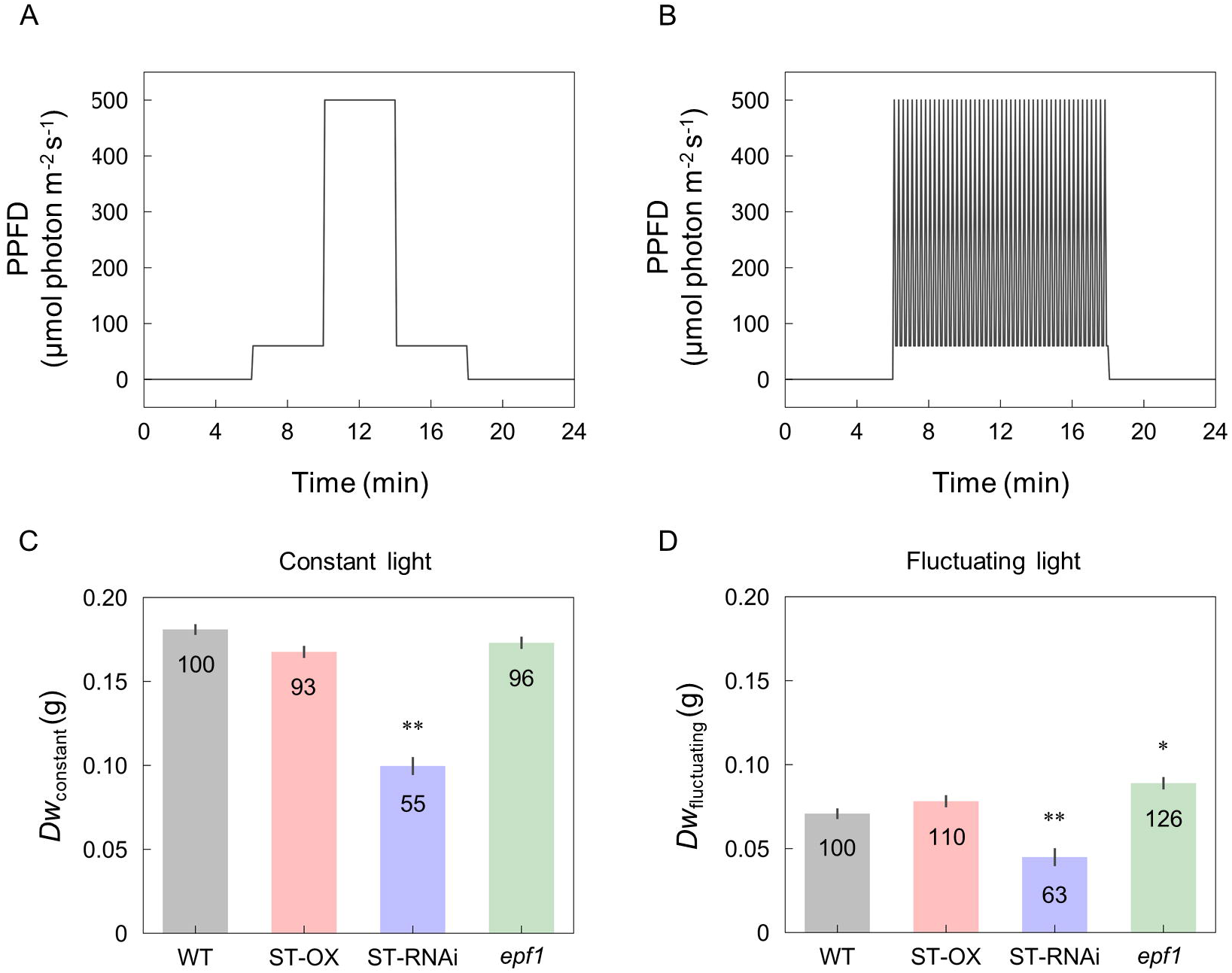
Biomass production under the constant and fluctuating light condition. Dry weight of the above ground biomass was evaluated in the four lines of Arabidopsis under (C) the constant (*DW*_constant_) and (D) fluctuating (*DW*_fluctuating_) light condition as described in (A) and (B). Vertical bars indicate the standard error (n = 4). * and ** indicate significant differences in each parameter between WT and each transgenic line at *p* < 0.05 and 0.01, respectively, according to Dunnett’s test. The values in each column represent the relative value of each line to WT.

## Discussion

Stomata play a significant role in the regulation of gas exchange between the outside and inside of the leaf. However, how the *SD* change affects photosynthetic and growth characteristics in plants has been controversial, and the effect of *SD* change on photosynthesis and growth can vary depending on the plant species or environmental conditions. Previously, we reported that higher *SD* resulted in the enhancement of *gs* and *A* in Arabidopsis under constant and saturated light conditions (Tanaka et al., 2013). Lawson and Blatt (2014) suggested that with higher *SD*, it would be instructive to determine biomass productivity under fluctuating light, although only a few studies investigated the relationship between *SD* and photosynthetic or growth characteristics under that condition (Drake et al., 2013; Papanatsiou et al., 2016; Schuler et al., 2018). Here, we attempted to examine how the change in *SD* affects *gs* and *A* dynamics, biomass production, and water use in Arabidopsis under fluctuating light.

In this study, the four Arabidopsis lines with contrasting *SDs* showed significant differences in the light response of *gs* in the steady and non-steady state. *SD* differences had no significant effect on the response of *gs* to the light intensity change from low to high in the present (Fig. 4A and B) and previous studies (Papanatsiou et al., 2016; Schuler et al., 2018). Contrastingly, ST-OX and ST-RNAi showed *gs* responses slower or faster, respectively, than that of WT when pre-adapted to the dark condition (Fig. 3A–C). This suggests that the relationship between *SD* and *gs* dynamics can depend on the light condition history of the leaf. The different responses of *gs* could be attributable to the difference in the size, density, and patterning of stomata. Drake *et al*. (2013) reported that smaller stomata respond the fluctuating light faster than larger stomata among several species of the genus *Banksia*. On the contrary, smaller stomata resulted in the slower response of *gs* to fluctuating light in the genus *Oryza* (Zhang et al., 2019). In this study, the variation in stomatal size evaluated as *L*_g_ did not correspond to that of *gs* response (Fig. 1 and 3), indicating that the stomatal size would have a minor effect on the dynamics of *gs* in Arabidopsis under fluctuating light.

The stomatal opening is regulated by at least three key components, blue-light receptor phototropin, plasma membrane H^+^-ATPase, and plasma membrane inward rectifying K^+^ channels in the guard cell (Inoue and Kinoshita, 2017). The activation of H^+^-ATPase induced by blue light as the initial signal facilitates K^+^ uptake through the inward rectifying K^+^ channel to increase the turgor pressure of guard cells, resulting in the stomatal opening. In addition, stomatal opening dynamics depend on the water status in the plant (Lawson and Blatt, 2014). With more stomata, higher metabolic cost and water uptake would be required for the movement. The gas-exchange and theoretical-modeling analysis indicated that the stomatal clustering decreased the maximum value of *gs* and *A* under the steady state because of the misplacement of stomatal pores over mesophyll cells (Dow. *et al*., 2014; Lehmann and Or, 2015). It was also shown that clustering suppressed stomatal movement owing to the decreased capacity of the K^+^ flux and K^+^ accumulation in the guard cells (Papanatsiou et al., 2016). Additionally, *gs* response to fluctuating light in *Begonia* species with clustered stomata was slower than that in those without clustered stomata (Papanatsiou et al., 2017). In this study, ST-OX, with the highest *SD* and clustering rate, showed the slowest *gs* response among Arabidopsis lines, while ST-RNAi, with the lowest *SD* and no clustered stomata, showed the fastest response (Fig. 1 and 3). *epf1*, with a moderate increase in *SD* and clustering rate, showed *gs* response similar to that of WT. These results suggest that the drastic change in stomatal density and patterning rather than size can largely affect *gs* response to fluctuating light owing to the change in metabolic cost and water uptake for stomatal opening and the opening speed of single stoma.

Stomatal patchiness, the spatially or temporally-heterogeneous distribution of stomatal apertures on the leaf, can occur in response to light fluctuation and affects the kinetics of the stomatal response (Mott and Buckley, 2000). Patchiness has been observed in many studies under water-limited conditions such as low humidity or soil water potential (Terashima, 1992). ST-OX showed higher *E* under the steady and non-steady state than WT (Fig. 2), leading to increased water loss during growth. This might induce stomatal patchiness more severely in ST-OX, which would require longer time to reach the full induction of *gs* than WT. This consideration can be partly supported by the fact that ST-RNAi, with the lowest *E* and highest *WUE*, showed the fastest *gs* response to fluctuating light of all lines (Fig. 1–3).

In ST-OX, ST-RNAi, and *epf1*, *A* response to light intensity change from dark to high intensity was faster than that of WT (Fig. 3D–F). Photosynthetic induction is typically limited by three phases of the biochemical or diffusional processes; (1) activation of electron transport, (2) activation of the enzymes of the Calvin-Benson cycle, and (3) stomatal opening (Pearcy, 1990; Yamori, 2016). The significance of stomatal limitation to photosynthetic induction depends on the initial value of *gs* (Kirschbaum and Pearcy, 1988). Activation of the electron transport and enzymes of the Calvin-Benson cycle can be largely affected by CO_2_ concentration when the small portion of Rubisco is activated (Jackson et al., 1991; Kaiser et al., 2017; Urban et al., 2008). The variation of *gs* under dark or low light conditions corresponded to that in the speed of photosynthetic induction in several plant species (Kaiser et al., 2016; Soleh et al., 2017). Kaiser *et al*., (2016) reported that *t*_dark_*A*0.9_ correlated with initial *gs* when *gs* was less than 0.13 mol m^-2^ s^-1^. In this study, *t*_dark_*A*0.6_ correlated with *gs*_dark_ if *gs*_dark_ < 0.055 mol m^-2^ s^-1^, and it was constant regardless of *gs*_dark_ for *gs*_dark_ > 0.055 mol m^-2^ s^-1^ (Fig. 5B). *gs*_dark_ of WT, ST-OX, and *epf1* were 0.032, 0.118, and 0.085 mol m^-2^s^-1^, respectively, suggesting that the variation in *gs*_dark_ would cause the response difference of *A* (Fig. 5A and B). Contrastingly, ST-RNAi showed a slight difference in *gs*_dark_ from WT, but the faster response of *gs*, which contributed to higher *gs* and *C*_i_ than those of WT in the initial induction phase (Fig. 2, 3, and 5). Therefore, higher *SD* resulted in higher initial value of *gs* while lower *SD* resulted in faster *gs* response, which would contribute to faster photosynthetic induction owing to the rapid activation of RuBP regeneration and carboxylation in the Calvin-Benson cycle.

After the change in light intensity from low to high, the response of *A* had no relation to initial *gs* because *gs*_low_ was over 0.1 mol m^-2^ s^-1^ in all lines (Fig. 5D–F). Thus, *gs* would not be a major limitation factor to photosynthetic induction. *t*_low_*A*0.6_ in the three transgenic lines was shorter than that in WT (*p* < 0.05), and *t*_low_*A*0.9_ in ST-OX was 52.8% shorter than that of WT (*p* < 0.1), suggesting that the activation of RuBP regeneration and carboxylation in the Calvin-Benson cycle in the transgenic lines might be faster than that in WT. Metabolic limitation to photosynthetic induction will be examined among Arabidopsis lines with contrasting *SD*s in a future study.

*DW*_fluctuating_ was much lower than *DW*_constant_ in all Arabidopsis lines, although the total amount of light intensity exposed to the plants was equal between both light conditions. This difference would be caused by the loss of carbon gain owing to photosynthetic induction under fluctuating light condition (Fig. 7). While there was no variation in *A* between WT and ST-RNAi in the steady state (Fig. S1), *DW*_constant_ in ST-RNAi was significantly lower than that in WT as reported in Tanaka *et al*. (2013) (Fig. 7C). In Arabidopsis, a near linear relationship was reported between leaf area and plant biomass during the vegetative stage (Weraduwage et al., 2015). Furthermore, Tanaka *et al*. (2013) showed a significant decrease in leaf area of ST-RNAi, suggesting that the silencing of *STOMAGEN*/*EPLF9* would decrease biomass production owing to the significant decrease in leaf area. Contrastingly, transgenic lines with lower *SD* exhibited improved performance in growth under drought stress owing to high *WUE* in several plant species (Caine et al., 2018; Wang et al., 2016; Yoo et al., 2010). Additionally, the rapid movement of stomata can be important for plants to achieve high *WUE* (Qu et al., 2016). In this study, compared with WT, ST-RNAi, with lower *SD*, showed faster *gs* response, similar amount of carbon gain, lower water loss, and lower biomass production loss caused by photosynthetic induction under fluctuating light condition (Fig. 3, 6, and 7). It is indicated that lower *SD* would be advantageous for plant growth under water-limited and fluctuating light condition.

*DW*_constant_ in ST-OX was slightly lower than that in WT, although *A*_max_ was significantly or slightly higher in Tanaka *et al*. (2013) and this study, respectively (Figs. 7 and S1). The increase in water loss and metabolic cost of stomatal movements would have a negative effect on biomass production in ST-OX under constant light (Tanaka et al., 2013). Although these penalties resulting from the drastic increase in *SD* might have negative effects on carbon gain, *DW*_fluctuating_ in ST-OX was 10.5% higher than that in WT with no significant variation. Moreover, biomass production in *epf1*, with moderate increase in *SD*, was significantly higher than that in WT under fluctuating light, but there was no difference between these two lines under constant light (Fig. 7). It is possible that a moderate increase in *SD* could achieve more efficient carbon gain attributable to the high capacity and fast response of *A* in Arabidopsis under fluctuating light (Fig. 1–6), while it would cause small penalties on water loss and metabolic cost for stomatal movement. Overall, higher *SD* can be beneficial to improve biomass production in plants under fluctuating light conditions.

### Conclusion

Under fluctuating light, there was a significant variation in the photosynthetic and growth characteristics among Arabidopsis lines contrasting in the stomatal density, size, and patterning. Lower *SD* resulted in faster *A* response owing to the faster *gs* response and higher *WUE* without the decrease in *A*. This suggests that lower *SD* would be advantageous for plant growth under water-limited and fluctuating-light conditions. Higher *SD* also resulted in faster *A* response to fluctuating light owing to the higher initial value of *gs*. *epf1*, with a moderate increase in *SD*, achieved more efficient carbon gain under fluctuating light attributable to the high capacity and fast response of *A*, which contributed to higher biomass production than that in WT. This study suggests that higher *SD* can be beneficial to improve biomass production in plants under field conditions.

## Materials and Methods

### Plant materials and cultivation for the gas exchange and biomass analyses

Columbia-0 (CS60000) of *Arabidopsis thaliana* (L.) Heynh, was used as a wild-type line (WT). In addition, we used three transgenic lines, *STOMAGEN/EPFL9* overexpressing (ST-OX) and artificial microRNA (amiRNA)-mediated silencing (ST-RNAi) lines, and an *EPF1* knockout line (*epf1*), which were developed by Sugano *et al*. (2010). The six plants per line were sown and grown on the soil at an air temperature of 22°C and a photosynthetic photon flux density (PPFD) of 100 µmol photon m^-2^ s^-1^ for the gas exchange analysis. The day/night length was set to 8/16 h. For the biomass analysis, plants were sown and grown on the soil at an air temperature of 22°C and a PPFD of 120 µmol photon m^-2^ s^-1^ for 24 days after sowing with the day/night length of 8/16 h. Subsequently, four plants per line were subjected to constant and fluctuating light conditions, for 20 days with a day/night cycle of 12/12 h. During daytime, the light intensity in the constant light condition was changed from a PPFD of 60 μmol m^-2^ s^-1^ for 4 h to 500 μmol m^-2^ s^-1^ for 4h, followed by 60 μmol m^-2^ s^-1^ for 4h, while a PPFD of 60 μmol m^-2^ s^-1^ for 10 min after 500 μmol m^-2^ s^-1^ for 5 min was repeated for 12 h in the fluctuating light condition (Fig. 7A and B). Plants were exposed to the same total amount of light intensity per day under both light conditions. Dry weight of above ground biomass grown under each light condition was evaluated at 44 days after sowing.

### Evaluation of stomatal density, size, and clustering

*SD*, guard cell length (*L_g_*), and stomatal clustering were evaluated in the leaves of the six plants per line at the same growth stage as the gas exchange measurements were conducted. A section of the leaf (5 × 5 mm) was excised and immediately fixed in the solution (Ethanol : acetic acid = 9:1, v/v) overnight. The fixed tissues were cleared in chloral hydrate solution (chloral hydrate : glycerol : water = 8:1:2, w/v/v) overnight. The cleared tissues were stained with safranin-O solution (200 μg ml^-1^) for 30 min to 1 h. The abaxial side of the leaves was observed at a 200× magnification using an optical microscope and digital images were obtained (CX31 and DP21, Olympus, Tokyo, Japan). We used imaging analysis software, ImageJ (NIH, Bethesda, MD, USA) to assess the stomatal number and guard cell length from the images. The percentage of clustered stomata to total number was measured for each clustering category.

### Gas exchange measurements

Gas exchange measurements were conducted using a portable gas-exchange system LI-6400 (*LI-COR*, Lincoln, NE, USA). All plants were kept in the dark overnight before and during the measurements. In the leaf chamber, we set flow rate at 300 µmol s^-1^, CO_2_ concentration at 400 µmol mol^-1^, and air temperature at 25°C. After the leaf was clamped in the chamber, light intensity was kept at dark condition for the initial 10 min and, subsequently, under a series of 500, 50, 500, 50, 500, 50, and 500 μmol photon m^-2^ s^-1^ for 120, 1, 20, 5, 20, 30, and 30 min as described in Fig. 2A. *A*, *gs*, intercellular CO_2_ concentration (*C*_i_), and *E* were recorded every 10 s during the measurements. *WUE* was calculated as the ratio of *A* to *E*. Gas exchange measurements were conducted with three plants per line during 68 to 73 days after sowing.

### Data processing and statistical analysis

The response of *A* and *gs* was evaluated at 500 μmol photon m^-2^ s^-1^ for 120 min after the dark period and at 500 μmol photon m^-2^ s^-1^ for 30 min after the illumination with 50 μmol photon m^-2^ s^-1^ for 30 min, based on *A*_induction_ and *gs*_induction_ defined as the following equations:

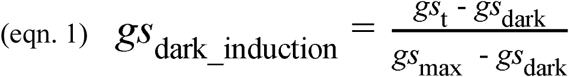

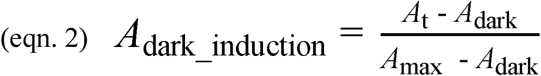

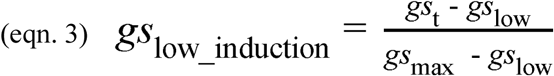

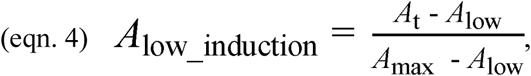

where *A*_dark_ and *gs*_dark_ represent steady-state values under the dark condition, *A*_low_ and *gs*_low_ are steady-state values under a PPFD of 50 μmol photon m^-2^ s^-1^, *A*_max_ and *gs*_max_ are maximum values under a PPFD of 500 μmol photon m^-2^ s^-1^, and *A*_t_ and *gs*_t_ represent *A* and *gs* at a given time under illumination. We evaluated the differences in the time when *A*_induction_ and *gs*_induction_ reached 0.6 and 0.9 after the light intensity change from dark to high (*t*_dark*_gs*0.6_, *t*_dark*_gs*0.9_, *t*_dark*_A*0.6_, and *t*_dark*_A*0.9_) or from low to high (*t*_low*_gs*0.6_, *t*_low*_gs*0.9_, *t*_low*_A*0.6_, and *t*_low*_A*0.9_) between WT and each transgenic line. To compare the photosynthetic characteristics under the steady state, we evaluated *A*, *g*s, *C*_i_, and *E* in the four lines when *A* reached a maximum value throughout the measurement.

The cumulative CO_2_ assimilation rate (*CAR*) and transpiration rate (*CER*) under fluctuating light were calculated by summing *A* and *E* after the initial dark period. An integrated *WUE* (*WUE*_i_) was calculated as the ratio of *CAR* to *CER*. Assuming the absence of induction response of *A* to fluctuating light, a theoretical maximum *CAR* (*CAR*_t_) was defined by the following equation:

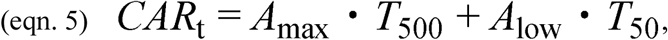

where *T*_500_ and *T*_50_ are the seconds for which the light intensity was maintained at 500 and 50 μ mol photon m^-2^ s^-1^, respectively. The potential loss rate of carbon gain caused by photosynthetic induction was defined by the following equation:

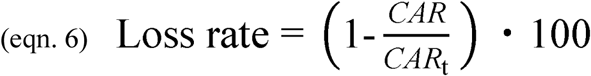

The variation in stomatal size and all the parameters of photosynthetic and growth characteristics were compared between WT and each transgenic line by a Dunnett’s test. Steel test was applied to evaluate *SD* variation between WT and each transgenic line because the distribution of values was extremely different among the lines. Statistical analysis was conducted using R software version 3. 6. 1 (R Foundation for Statistical Computing, Vienna, Austria).

## Acknowledgements

We are grateful to Mr. S. Shimadzu for help in the biomass analysis.

## References

Adachi, S., Tanaka, Y., Miyagi, A., Kashima, M., Tezuka, A., Toya, Y., et al. (2019) High-yielding rice Takanari has superior photosynthetic response under fluctuating light to a commercial rice Koshihikari. J Exp Bot. 70: 5287–5297.

von Caemmerer, S., and Evans, J.R. (2010) Enhancing C3 Photosynthesis. Plant Physiol. 154: 589–592.

Caine, R.S., Yin, X., Sloan, J., Harrison, E.L., Mohammed, U., Fulton, T., et al. (2018) Rice with reduced stomatal density conserves water and has improved drought tolerance under future climate conditions. New Phytol. 221: 371–384.

Carmo-Silva, A.E., and Salvucci, M.E. (2013) The Regulatory Properties of Rubisco Activase Differ among Species and Affect Photosynthetic Induction during Light Transitions. Plant Physiol. 161: 1645–1655.

Doheny-Adams, T., Hunt, L., Franks, P.J., Beerling, D.J., and Gray, J.E. (2012) Genetic manipulation of stomatal density influences stomatal size, plant growth and tolerance to restricted water supply across a growth carbon dioxide gradient. Philos Trans R Soc B Biol Sci. 367: 547–555.

Dow, G.J., Berry, J.A., Bergmann, D.C. (2014) The physiological importance of developmental mechanisms that enforce proper stomatal spacing in Arabidopsis thaliana. New Phytol. 201: 1205–1217.

Drake, P.L., Froend, R.H., and Franks, P.J. (2013) Smaller, faster stomata: Scaling of stomatal size, rate of response, and stomatal conductance. J Exp Bot. 64: 495–505.

Farquhar, G.D., and Sharkey, T.D. (1982) Stomatal Conductance and Photosynthesis. Annu Rev Plant Physiol. 33: 317–345.

Franks, P.J., and Beerling, D.J. (2009) CO2-forced evolution of plant gas exchange capacity and water-use efficiency over the Phanerozoic. Geobiology. 7: 227–236.

Hara, K., Kajita, R., Torii, K.U., Bergmann, D.C., and Kakimoto, T. (2007) The secretory peptide gene EPF1. Genes Dev. 1720–1725.

Hashimoto-Sugimoto, M., Higaki, T., Yaeno, T., Nagami, A., Irie, M., Fujimi, M., et al. (2013) A Munc13-like protein in Arabidopsis mediates H+-ATPase translocation that is essential for stomatal responses. Nat Commun. 4: 1–7.

Inoue, S., and Kinoshita, T. (2017) Blue Light Regulation of Stomatal Opening and the Plasma Membrane H ^+^ -ATPase. Plant Physiol. 174: 531–538.

Jackson, R.B., Woodrow, I.E., and Mott, K.A. (1991) Nonsteady-state photosynthesis following an increase in photon flux density (PFD). Plant Physiol. 95: 498–503.

Kaiser, E., Kromdijk, J., Harbinson, J., Heuvelink, E., and Marcelis, L.F.M. (2017) Photosynthetic induction and its diffusional, carboxylation and electron transport processes as affected by CO 2 partial pressure, temperature, air humidity and blue irradiance. Ann Bot. 119: 191–205.

Kaiser, E., Morales, A., Harbinson, J., Heuvelink, E., Prinzenberg, A.E., and Marcelis, L.F.M. (2016) Metabolic and diffusional limitations of photosynthesis in fluctuating irradiance in Arabidopsis thaliana. Sci Rep. 6: 1–13.

Kirschbaum, M.U.F., and Pearcy, R.W. (1988) Gas Exchange Analysis of the Fast Phase of Photosynthetic Induction in Alocasia macrorrhiza. Plant Physiol. 87: 818–821.

Lawson, T., and Blatt, M.R. (2014) Stomatal size, speed, and responsiveness impact on photosynthesis and water use efficiency. Plant Physiol. 164: 1556–70.

Lee, J.S., Hnilova, M., Maes, M., Lin, Y.C.L., Putarjunan, A., Han, S.K., et al. (2015) Competitive binding of antagonistic peptides fine-tunes stomatal patterning. Nature. 522: 439–443.

Lehmann, Peter; Or, D. (2015) Supporting Information - Effects of stomatal clustering on leaf gas exchange. New Phytol. 207: 1015–1025.

Mcadam, S.A.M., and Brodribb, T.J. (2012) Stomatal innovation and the rise of seed plants. Ecol Lett. 15: 1–8.

Mott, K.A., and Buckley, T.N. (2000). Patchy stomatal conductance: Emergent collective behaviour of stomata. Trends Plant Sci. 5: 258–262.

Papanatsiou, M., Amtmann, A., and Blatt, M.R. (2016) Stomatal Spacing Safeguards Stomatal Dynamics by Facilitating Guard Cell Ion Transport Independent of the Epidermal Solute Reservoir. Plant Physiol. 172: 254–263.

Papanatsiou, M., Amtmann, A., and Blatt, M.R. (2017) Stomatal clustering in Begonia associates with the kinetics of leaf gaseous exchange and influences water use efficiency. J Exp Bot. 68: 2309–2315.

Papanatsiou, M., Petersen, J., Henderson, L., Wang, Y., Christie, J.M., and Blatt, M.R. (2019) Optogenetic manipulation of stomatal kinetics improves carbon assimilation, water use, and growth. Science. 363: 1456–1459.

Qi, X., and Torii, K.U. (2018) Hormonal and environmental signals guiding stomatal development. BMC Biol. 16: 1–11.

Qu, M., Hamdani, S., Li, W., Wang, S., Tang, J., Chen, Z., et al. (2016) Rapid stomatal response to fluctuating light: an under-explored mechanism to improve drought tolerance in rice. Funct Plant Biol. 43: 727.

Sakoda, K., Tanaka, Y., Long, S.P., and Shiraiwa, T. (2016) Genetic and physiological diversity in the leaf photosynthetic capacity of soybean. Crop Sci. 56: 2731–2741.

Schlüter, U., Muschak, M., Berger, D., and Altmann, T. (2003) Photosynthetic performance of an Arabidopsis mutant with elevated stomatal density (*sdd1-1*) under different light regimes. J Exp Bot. 54: 867–874.

Schuler, M.L., Sedelnikova, O. V., Walker, B.J., Westhoff, P., and Langdale, J.A. (2018) SHORTROOT-mediated increase in stomatal density has no impact on photosynthetic efficiency. Plant Physiol. 176: pp.01005.2017.

Shimadzu, S., Seo, M., Terashima, I., and Yamori, W. (2019) Whole Irradiated Plant Leaves Showed Faster Photosynthetic Induction Than Individually Irradiated Leaves via Improved Stomatal Opening. Front Plant Sci. 10: 1–10.

Soleh, M.A., Tanaka, Y., Kim, S.Y., Huber, S.C., Sakoda, K., and Shiraiwa, T. (2017) Identification of large variation in the photosynthetic induction response among 37 soybean [Glycine max (L.) Merr.] genotypes that is not correlated with steady-state photosynthetic capacity. Photosynth Res. 131: 305–315.

Soleh, M.A., Tanaka, Y., Nomoto, Y., Iwahashi, Y., Nakashima, K., Fukuda, Y., et al. (2016) Factors underlying genotypic differences in the induction of photosynthesis in soybean [Glycine max (L.) Merr.]. Plant Cell Environ. 39: 685–693.

Sugano, S.S., Shimada, T., Imai, Y., Okawa, K., Tamai, A., Mori, M., et al. (2010) Stomagen positively regulates stomatal density in Arabidopsis. Nature. 463: 241–244.

Tanaka, Y., Adachi, S., and Yamori, W. (2019) Natural genetic variation of the photosynthetic induction response to fluctuating light environment. Curr. Opin. Plant Biol. 49: 52–59

Tanaka, Y., Sugano, S.S., Shimada, T., and Hara-Nishimura, I. (2013) Enhancement of leaf photosynthetic capacity through increased stomatal density in Arabidopsis. New Phytol. 198: 757–764.

Taylor, S.H., and Long, S.P. (2017) Slow induction of photosynthesis on shade to sun transitions in wheat may cost at least 21% of productivity. Philos Trans R Soc B Biol Sci. 372.

Terashima, I. (1992) Anatomy of non-uniform leaf photosynthesis. Photosynth Res. 31: 195–212.

Urban, O., Šprtová, M., Košvancová, M., Tomášková, I., Lichtenthaler, H.K., and Marek, M. V. (2008) Comparison of photosynthetic induction and transient limitations during the induction phase in young and mature leaves from three poplar clones. Tree Physiol. 28: 1189–1197.

Wang, C., Liu, S., Dong, Y., Zhao, Y., Geng, A., Xia, X., et al. (2016) *PdEPF1* regulates water-use efficiency and drought tolerance by modulating stomatal density in poplar. Plant Biotechnol J. 14: 849–860.

Weraduwage, S.M., Chen, J., Anozie, F.C., Morales, A., Weise, S.E., and Sharkey, T.D. (2015) The relationship between leaf area growth and biomass accumulation in Arabidopsis thaliana. Front Plant Sci. 6: 1–21.

Wong, S.C., Cowan, I.R., and Farquhar, G.D. (1979) Stomatal conductance correlates with photosynthetic capacity. Nature. 282: 424–426.

Yamori, W. (2016) Photosynthetic response to fluctuating environments and photoprotective strategies under abiotic stress. J Plant Res. 129: 379–395.

Yamori, W., Kondo, E., Sugiura, D., Terashima, I., Suzuki, Y., and Makino, A. (2016) Enhanced leaf photosynthesis as a target to increase grain yield: Insights from transgenic rice lines with variable Rieske FeS protein content in the cytochrome b6/f complex. Plant Cell Environ. 39: 80–87.

Yamori, W., Masumoto, C., Fukayama, H., and Makino, A. (2012) Rubisco activase is a key regulator of non-steady-state photosynthesis at any leaf temperature and, to a lesser extent, of steady-state photosynthesis at high temperature. Plant J. 71: 871–880.

Yoo, C.Y., Pence, H.E., Jin, J.B., Miura, K., Gosney, M.J., Hasegawa, P.M., et al. (2010) The Arabidopsis GTL1 transcription factor regulates water use efficiency and drought tolerance by modulating stomatal density via transrepression of *SDD1*. Plant Cell. 22: 4128–4141.

Zhang, Q., Peng, S., and Li, Y. (2019) Increase rate of light-induced stomatal conductance is related to stomatal size in the genus Oryza. J Exp Bot. 70: 5259–5269.

