## Supplementary Figure 1 for "Stomatal density affects gas diffusion and CO_2_ assimilation dynamics in Arabidopsis under fluctuating light"

Stomatal density (*SD*) is closely associated with photosynthetic and growth characteristics in plants. In the field, light intensity can fluctuate drastically throughout a day. The objective of the present study is to examine how the change in *SD* affects stomatal conductance (*gs*) and $CO_2$ assimilation rate (*A*) dynamics, biomass production, and water use under fluctuating light. Here, we compared the photosynthetic and growth characteristics under constant and fluctuating light among four lines of *Arabidopsis thaliana* (L.): a wild-type (WT), a *STOMAGEN/EPFL9*-overexpressing line, *STOMAGEN/EPFL9*-silencing line, and an *EPIDERMAL PATTERNING FACTOR 1* knockout line (*epf1*). Lower *SD* resulted in faster response of *A* owing to the faster response of *gs* to fluctuating light and higher water use efficiency without decreasing *A*. Higher *SD* resulted in a faster response of *A* because of the higher initial *gs*. *epf1*, with a moderate increase in *SD*, showed the larger carbon gain, attributable to the high capacity and fast response of *A*, yielding higher biomass production than WT under fluctuating light. The present study suggests that higher *SD* can be beneficial to improve biomass production in the plant under field conditions.

Fig. S1. Photosynthetic characteristics under the steady state

(A) stomatal conductance (*gs*), (B) intercellular $CO_2$ concentration ($C_i$), (C) $CO_2$ assimilation rate (*A*), (D) transpiration rate (*E*) and (E) water use efficiency (*WUE*) were measured on the fully expanded leaf in the four lines of Arabidopsis. The gas exchange measurements were conducted at a $CO_2$ concentration of 400 ppm, PPFD of 500 µmol $m^{-2}$ $s^{-1}$, and air temperature of 25 °C. The vertical bars indicate the standard error (n = 3). * indicates the significant variation between WT and each transgenic line at $p < 0.05$, according to the Dunnett's test. The values in each column represent the relative value of each line to WT.

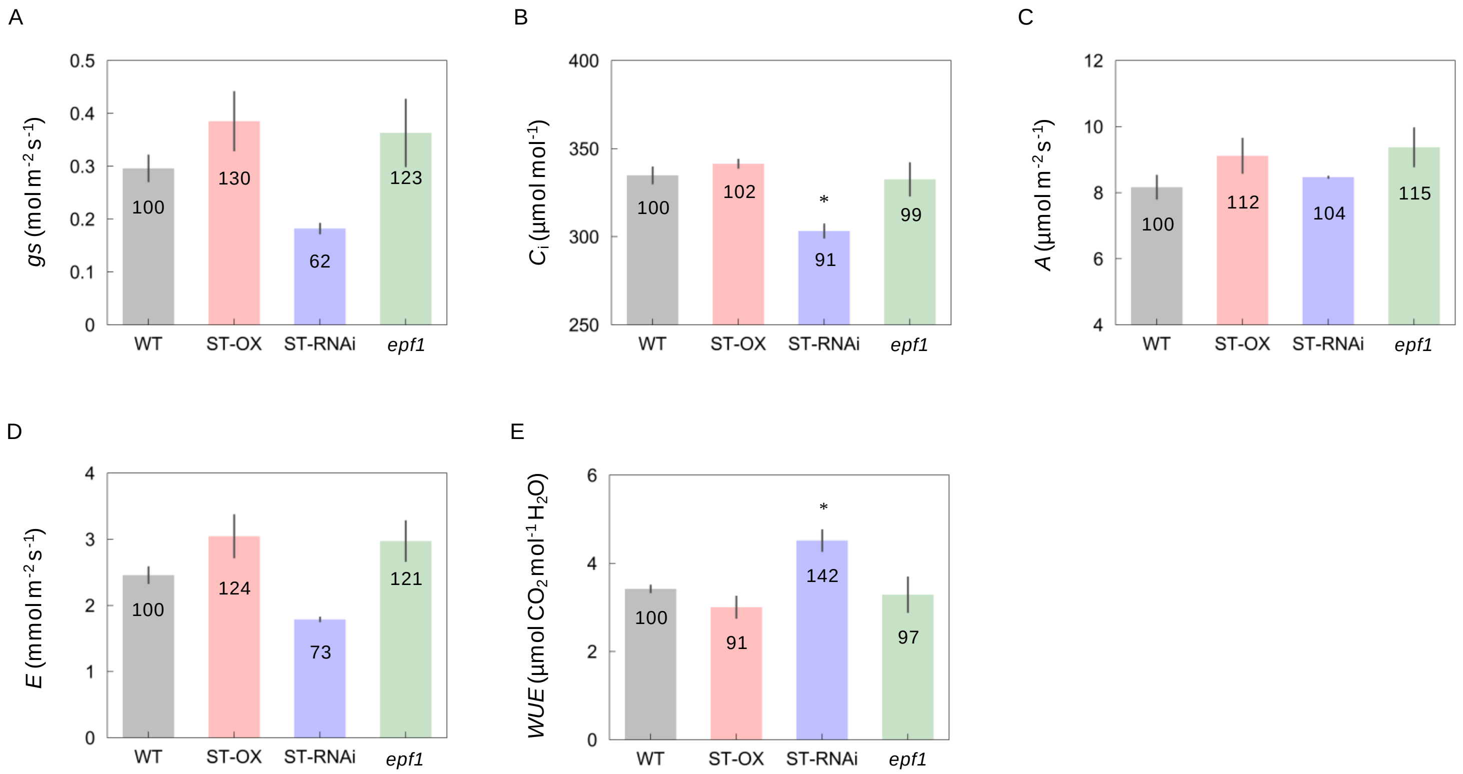
